# Effective polyploidy causes phenotypic delay and influences bacterial evolvability

**DOI:** 10.1101/226654

**Authors:** Lei Sun, Helen K. Alexander, Balazs Bogos, Daniel J. Kiviet, Martin Ackermann, Sebastian Bonhoeffer

## Abstract

Whether mutations in bacteria exhibit a noticeable delay before expressing their corresponding mutant phenotype was discussed intensively in the 1940s-50s, but the discussion eventually waned for lack of supportive evidence and perceived incompatibility with observed mutant distributions in fluctuation tests. Phenotypic delay in bacteria is widely assumed to be negligible, despite lack of direct evidence. Here we revisited the question using recombineering to introduce antibiotic resistance mutations into *E. coli* at defined time points and then tracking expression of the corresponding mutant phenotype over time. Contrary to previous assumptions, we found a substantial median phenotypic delay of 3-4 generations. We provided evidence that the primary source of this delay is multifork replication causing cells to be effectively polyploid, whereby wild-type gene copies transiently mask the phenotype of recessive mutant gene copies in the same cell. Using modeling and simulation methods, we explored the consequences of effective polyploidy for mutation rate estimation by fluctuation tests and sequencing-based methods. For recessive mutations, despite the substantial phenotypic delay, the *per-copy*or *per-genome* mutation rate is accurately estimated. However, the *per-cell* rate cannot be estimated by existing methods. Finally, with a mathematical model, we showed that effective polyploidy increases the frequency of costly recessive mutations in the standing genetic variation, and thus their potential contribution to evolutionary adaptation, while drastically reducing the chance that *de novo* recessive mutations can rescue populations facing a harsh environmental change such as antibiotic treatment. Overall, we have identified phenotypic delay and effective polyploidy as previously overlooked but essential components in bacterial evolvability, including antibiotic resistance evolution.

**Author summary:** What is the time delay between the occurrence of a genetic mutation in a bacterial cell and manifestation of its phenotypic effect? We show that antibiotic resistance mutations in *E.coli* show a remarkably long phenotypic delay of 3-4 bacterial generations. The primary underlying mechanism of this delay is effective polyploidy. In a polyploid cell with multiple chromosomes, once a mutation arises on one of the chromosomes, the presence of non-mutated, wild-type gene copies on other chromosomes may mask the phenotype of the mutation. One implication of this finding is that conventional methods to determine the mutation rates of bacteria do not detect polyploidy and thus underestimate their potential for adaptation. More generally, the effect that a new mutation may become useful only in the “grand-children of the grand-children” suggests that pre-existing mutations are more important for surviving sudden environmental catastrophe.

## Introduction

As genetic mutations appear on the DNA, their effects must first transcend the RNA and protein levels before resulting in an altered phenotype. This so-called “phenotypic delay” in the expression of new mutations could have major implications for evolutionary adaptation, particularly if selection pressures change on a time scale that is short relative to this delay, as may be the case for selection by antibiotics. The duration of phenotypic delay is an old but nearly forgotten question in microbiology(1-3). Luria and Delbrück were interested in the delay as they expected it to affect the mutant distribution in the fluctuation test in their seminal work on the random nature of mutations(1). They argued that if a mutant clone expressed its phenotype after *G* generations, then phenotypic mutants should be observed in populations in groups of 2^G^. Frequent observations of single-mutant populations however suggested G≈0. They thus concluded that the phenotypic delay is negligible(1,3,4). This has remained the *modus operandi*(4), despite the fact that molecular cloning protocols imply a significant delay as they require a waiting time typically longer than a bacterial generation to express new genetic constructs(5).

To quantify the phenotypic delay more directly, the time point of occurrence of a mutation in a cell needs to be known, which has only become possible with modern methods of genetic engineering. Here we use a recombineering approach to introduce mutations in *E. coli* within a narrow time window and find a remarkable phenotypic delay of 3-4 generations for three antibiotic resistance mutations. We identify the underlying mechanism as effective polyploidy, which reconciles the long phenotypic delay with Luria and Delbrücks observations. Investigating the consequences of effective polyploidy and phenotypic delay, we find that mutation rate estimates need to be adjusted for ploidy. Moreover, resistance mutations that occur after exposure to antibiotics are much less likely to survive due to the multi-generational phenotypic delay, while pre-existing mutations become a much more important contributor to survival.

## Results

### Mutations in bacteria exhibit multi-generational phenotypic delay

To quantify phenotypic delay we introduced each of four mutations at a specified time-point in *E. coli* with an optimized recombineering protocol (Methods), in which an ssDNA-oligonucleotide carrying the point-specific mutation is transformed into bacteria by electroporation(6). The ssDNA then binds reverse-complementarily to its target on the bacterial genome as part of a lagging-strand in an open replication fork(6), thereby introducing the mutation. Three mutations (RifR, NalR, StrepR) confer antibiotic resistance(7); the fourth mutation (lac+) enables lactose prototrophy(8) (Table S1). After introduction of the mutations, the cells grew continuously without selection and were sampled over time. Sampled cells were subjected to “immediate” versus “postponed” selection to quantify, respectively, the frequencies of current phenotypic mutants and of genotypic mutants (that contain at least one mutant gene copy and eventually have some phenotypic mutant descendants), with their ratio called phenotypic penetrance (Fig. 1A). Phenotypic delay is quantified as the time in bacterial generations to reach 50% phenotypic penetrance.

**Fig. 1.**
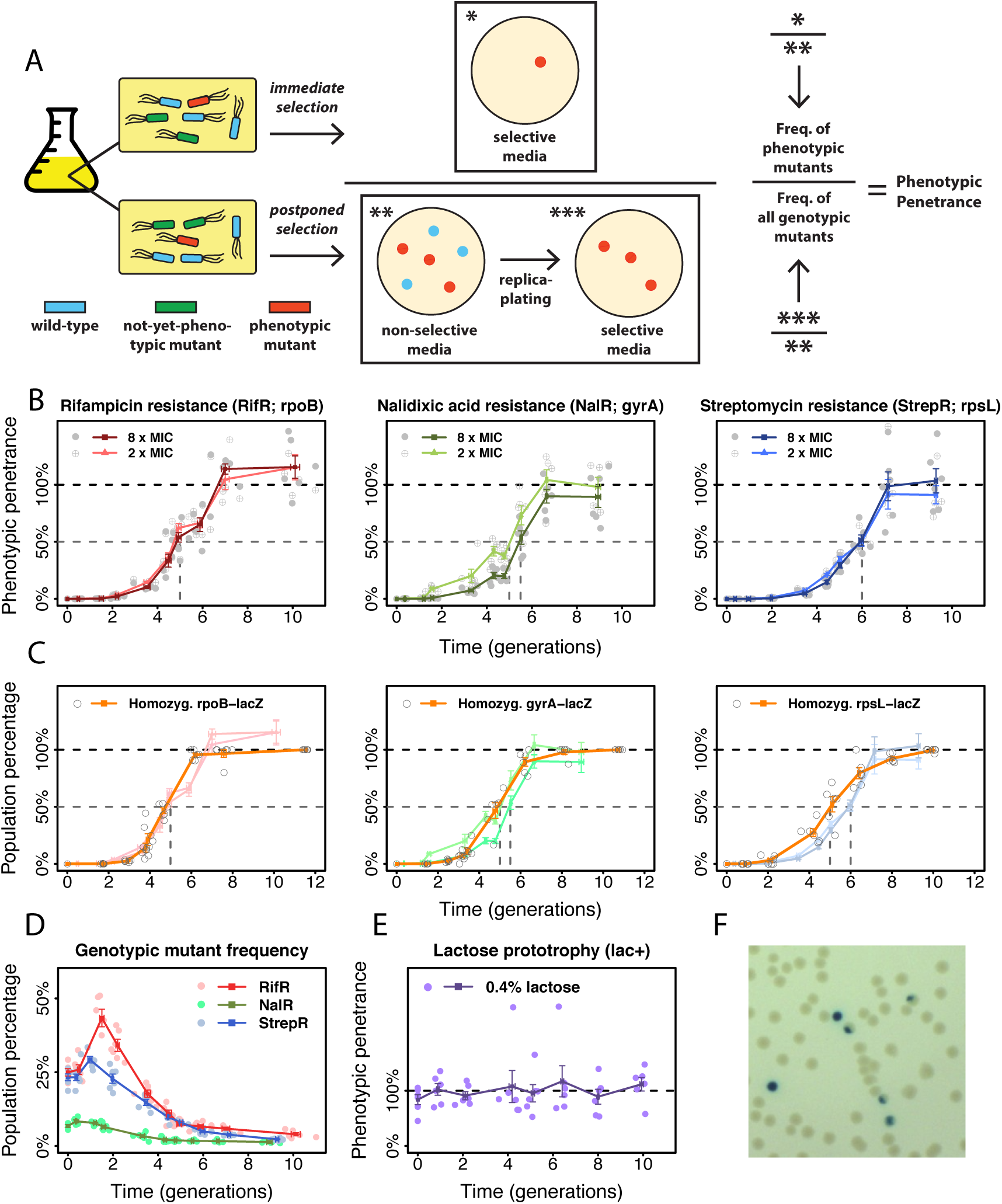
Phenotypic delay in *E. coli*. (A) Schematic illustration of the quantification of phenotypic penetrance. (B): phenotypic penetrance (mean ± SE; n = 6) over time for three antibiotic resistance mutations. Gray dashed lines: time at 50% phenotypic penetrance. (C): Frequency of homozygous mutants among all mutants (orange) for the three resistance mutations assessed by *lacZ*-reporter constructs (*rpoB*-*lacZ*, *gyrA*-*lacZ*, *rpsL*-*lacZ*), overlaid with their respective phenotypic penetrance. (D): Genotypic mutant frequency for the resistance mutations. (E): Phenotypic penetrance of the lactose prototrophy (*rpsL*-*lacZ*). (F): Colonies founded by homozygous (blue) and heterozygous (sectored) lac+ mutants.

Surprisingly, all three resistance mutations showed significant phenotypic delay at two selective concentrations of their respective antibiotic (Fig. 1B). Reaching 50% phenotypic penetrance required 5-6 generations of post-recombineering growth. The frequency of genotypic mutants increased over the first 1-2 generations but eventually declined (Fig. 1D). The transient increase may reflect the time window of introduction of the mutations. Discounting the first two generations, a phenotypic delay of 3-4 generations remains to be explained.

### Effective polyploidy is the primary source of phenotypic delay

Phenotypic delay could result from multiple factors. Firstly, it could arise from the gradual replacement of wild-type proteins by mutant proteins following mutagenesis. Time may be required for sufficient protein turnover before the mutant phenotype can manifest. Another possibility is that cells are effectively polyploid due to multifork replication(9,10). Recombineering incorporates the mutation into only one or some of the chromosomes starting from a single-strand mutant DNA(6), comparable to the occurrence of natural mutations. This yields effectively heterozygous cells that could produce both wild-type and mutant proteins from different chromosomes, which may prevent the onset of the mutant phenotype. Three generations could be the minimal time needed for a cell with one mutant copy out of eight chromosomes (comparable to previous estimates(9)) to produce the first homozygous mutant carrying only mutant alleles. Effective polyploidy is also compatible with the observed decline in genotypic mutant frequency (Fig. 1D), because heterozygous mutants produce both mutant and wild-type descendants, such that the frequency of cells carrying at least one mutant gene copy will decline until all cells are homozygous.

To quantify the contribution of effective polyploidy, we used a *lacZ* reporter assay to visualize heterozygous mutants. We constructed three reporter strains with a disrupted *lacZ* gene inserted close to each resistance target gene and restored it through recombineering (Table S2). Genotypic mutants were visualized by plating on indicator media where lac+ and lac cells become blue and white, respectively. Heterozygous mutants generate sectored colonies while homozygous mutants generate blue colonies, thus indicating the frequency of homozygous mutants amongst all genotypic mutants (Fig. 1F). Comparing the estimated proportion of homozygous reporter mutants with the corresponding phenotypic penetrance of the resistance mutation reveals that phenotypic delay can be fully explained by effective polyploidy for NalR at 2xMIC and for RifR (Fig. 1C). Homozygosity precedes phenotypic penetrance by about 0.5 generations for NalR at 8xMIC and one generation for StrepR, suggesting that here additional protein turnover may be involved.

These results also imply that these resistance mutations are genetically recessive to antibiotic sensitivity, which has also been described in previous studies based on co-expression assays(11,12). The recessive nature of these antibiotic resistance alleles stems from their molecular mechanism: when the antibiotic molecule binds to its target protein, the resulting complex is capable of damaging the cell even in small quantities, essentially acting as toxins. As a result, the gene dosage of wild-type targets is critical. For instance, nalidixic acid-bound gyrase proteins can introduce DNA double-stranded breaks(13). In particular, bacteria that overexpress gyrase become even more sensitive to quinolone antibiotics(14). Although the exact killing mechanism of streptomycin remains a subject of debate, it is generally accepted that streptomycin-bound ribosomes damage the cell via mistranslation(15). Finally, rifampicin-bound RNA polymerase appears to blockade the DNA, thereby preventing transcription even by drug-resistant RNA polymerases(12). Although dominant RifR mutations have also been described(16), the mutation we tested here appears to be recessive.

In the case of *lacZ* mutations, we scored the frequency of lac+ phenotypic mutants on lactose-limited medium. The ability to metabolize lactose is dominant to its inability(8). Since any cell containing a lac+ allele can metabolize lactose and eventually form a colony, phenotypic penetrance, as expected, was always at 100% (Fig. 1E), indicating that the observed phenotypic delay of resistance mutations is not an artifact of our protocol.

### Effective polyploidy causes asymmetrical inheritance of mutations

A further testable prediction of effective polyploidy is that inheritance of mutant alleles is asymmetrical: heterozygous mutants are expected to produce both wild-type and mutant offspring. For mutations with intermediate dominance, offspring progressing towards mutant homozygosity should show an increasingly mutant phenotype, while others show a transient phenotype as they inherit no mutant genes and their mutant proteins are diluted over subsequent divisions. Phenotypic delay would manifest as the time such mutations need to reach full phenotypic expression. To test this prediction, we repaired a disrupted *YFP* gene with recombineering, creating fluorescent mutants where the fluorescence intensity depends on the number of functional copies of this gene. We then tracked fluorescence as an intermediate-dominant phenotypic trait using single cell imaging. As expected we observed fully, transiently and non-fluorescent offspring lineages from recombineering-treated cells (Fig. 2A-C, Supplementary Movie), consistent with effective polyploidy. Furthermore, fluorescence in mutant lineages increased monotonically and reached maximal intensity almost two generations after forming homozygous mutants (Fig. 2C). This additional delay could be due to protein folding(17). In total we found 34 homozygous mutant lineages in 25 micro-colonies. Six micro-colonies spawned multiple, separate homozygous mutant lineages, thus corroborating previous findings that recombineering may modify multiple genome copies in one cell(18). Overall, a median of five generations was required to form homozygous mutants, consistent with our *lacZ* reporter assay results (Fig. 2D). These results provide direct visual support that effective polyploidy underlies phenotypic delay. A similar pattern has been observed previously in *E. coli* with fimbrial switching, a genetic modification that involves inversion of a promoter sequence on the bacterial genome(19).

**Fig. 2.**
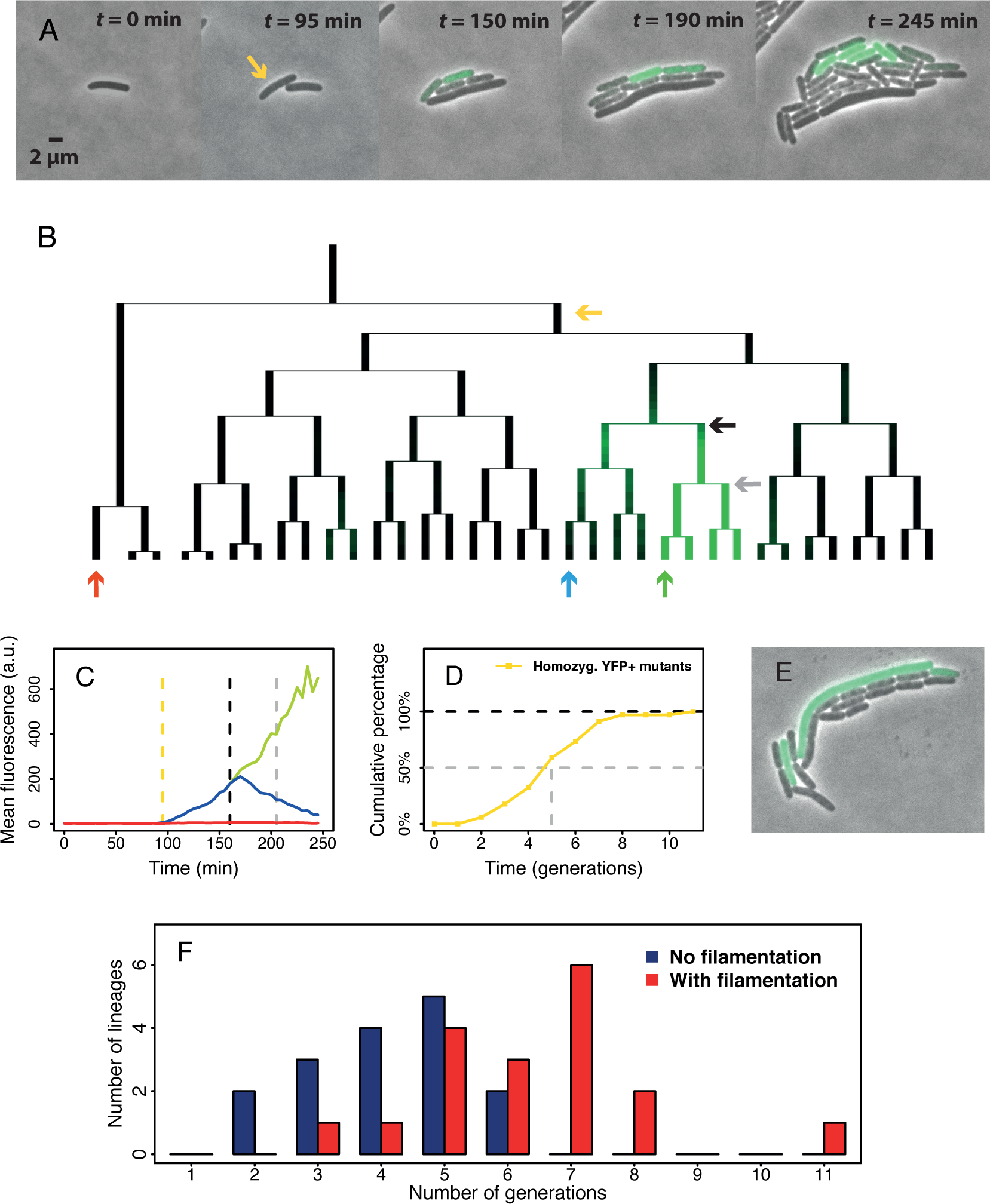
Single-cell analysis of fluorescent mutants. (A): Overlay of phase-contrast and fluorescence images showing a micro-colony containing fluorescent mutants (see also Supplementary Movie). Yellow arrow: the first cell showing significantly higher fluorescence than background in the given frame. Accounting for the time required for YFP protein folding and maturation, the ssDNA integration is estimated to have happened before the first division of the labeled cell. (B): Genealogy of the aforementioned micro-colony. The yellow arrow indicates the cell in (A) while the remaining arrows indicate three lineages in which fluorescence was quantified. (C): YFP expression history of three lineages in (B) showing fully, transiently and non-fluorescent phenotypes (green, blue and red). Yellow dashed line: onset of fluorescence. Black and grey dashed lines correspond to the black and grey arrows in (B). Black: emergence of the first homozygous mutant; grey: its first division. (D): Distribution of time to form 34 homozygous mutant lineages from 25 micro-colonies. The data is obtained by directly analyzing genealogies as in (B), and compiled from two separate experiments. (E): Photo of a micro-colony with one filamentous fluorescent cell. (F): The distribution of numberof generations to form homozygous mutant lineages sorted by the presence/absence of filamentation.

Electroporation, as used in the recombineering protocol, may cause bacteria to form filaments due to stress (20). Filamentation might exacerbate phenotypic delay by increasing the intracellular genome copy number. Our single-cell experiment revealed that filamentation was indeed frequent (Fig. 2E). By directly observing the cell shape and the time-point for onset of fluorescence, we estimated that 18 out of 34 homozygous mutant lineages incorporated the ssDNA mutation into a filamentous ancestor cell. Strikingly, lineages initiated by filamentous cells showed a distinctly different distribution of time to form the first homozygous mutant than non-filamentous lineages (Fig. 2F). Non-filamentous lineages showed a median of 4 generations until homozygosity, as would be required for cells that incorporated the mutation in 1 out of 16 DNA single-strands (i.e. 8 chromosomes) in less than one generation post-recombineering. In contrast, filamentous lineages showed a median of 6.5 generations, and all lineages requiring more than 6 generations were filamentous. In conclusion, both filamentous and non-filamentous cells exhibit a multi-generational phenotypic delay. Filamentation, however, can exacerbate phenotypic delay presumably by increasing the number of chromosome copies within a cell, explaining particularly long delays of five or more generations sometimes observed in our experiments.

### Effects of chromosomal location on mutagenesis and delay

Since phenotypic delay arises from effective polyploidy, one would expect genes further from the replication origin with lower ploidy than origin-proximal genes to show shorter phenotypic delay. However, we observed a similar phenotypic delay and distribution of time until homozygosity for all three tested genes (Fig. 1) despite different distances from the origin. Further analysis revealed a strong negative correlation between distance to origin and the initial frequency of genotypic mutants induced by recombineering (Fig. 3A-B). We hypothesized that these observations reflect the mechanism of mutagenesis by recombineering. Since open replication forks are required for recombineering, mutations could only be introduced during DNA replication(2 1). The probability of successful mutagenesis on at least one open chromosomal target increases with ploidy, which itself increases during DNA replication. Therefore we hypothesized that instead of mutating at low ploidy and thereby exhibiting shorter phenotypic delay, most of our observed origin-distal mutations were generated when their targets transiently reached higher ploidy either during normal DNA replication or due to cell filamentation. Thus, consistent with our observations so far, origin-distal mutations would show similar phenotypic delay as origin-proximal mutations, but reduced recombineering efficiency since origin-distal genes are less accessible for mutagenesis.

**Fig. 3.**
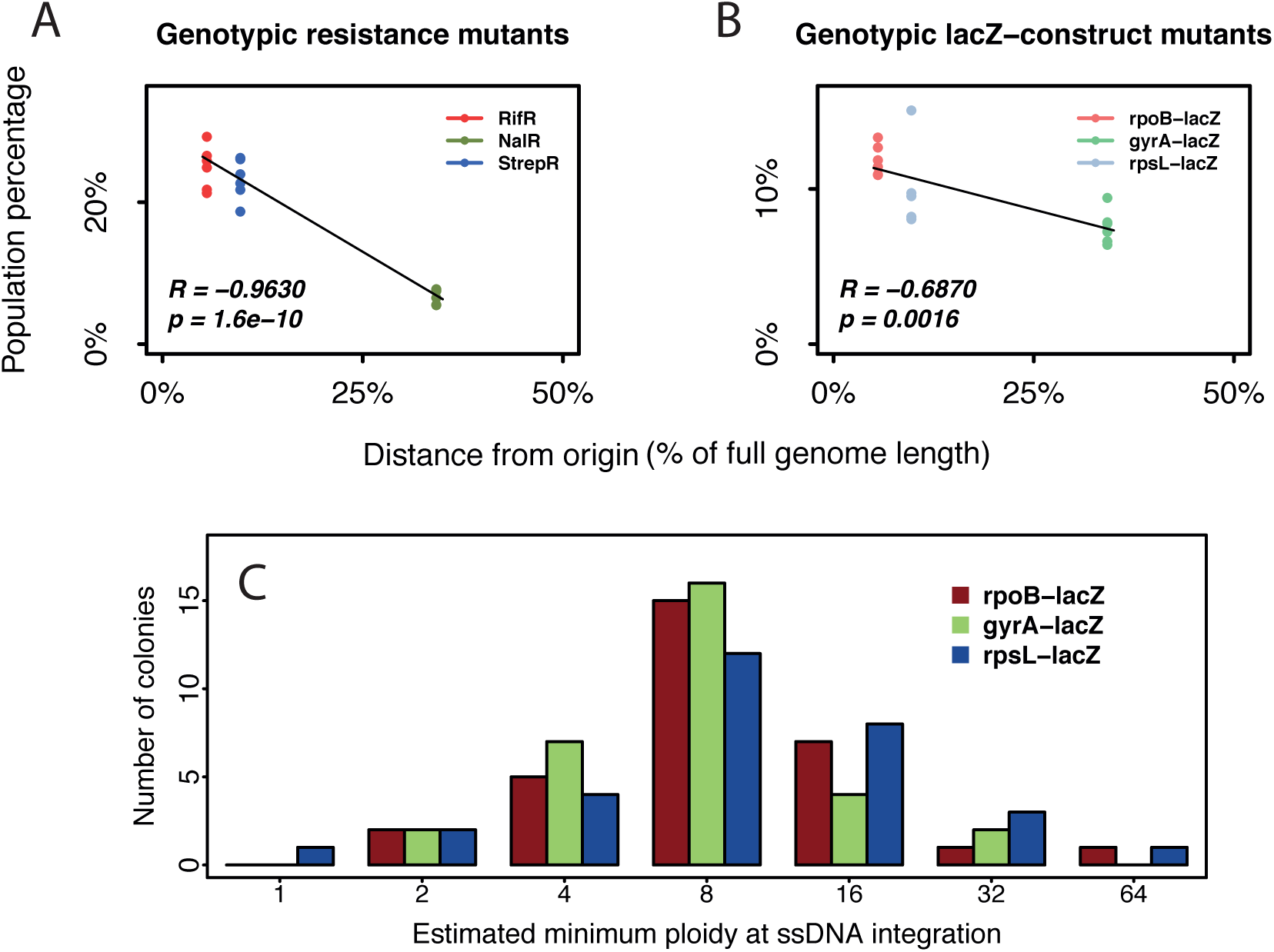
Chromosomal location effects on recombineering. Resistance target genes rpoB, gyrA and rpsL are at 5.5%, 34.2% and 9.7% genomic distance from the replication origin, respectively. Strong negative correlation (quantified by Pearsons correlation coefficient, R) exists between the distance from the origin and the initial frequency of genotypic mutants induced by recombineering, for both (A) the resistance mutations and (B) their *lacZ*-reporters. (C): The inferred minimal ploidy from the cell physically treated by recombineering at the time of mutant ssDNA integration shows a similar distribution for all constructs, regardless of their distance from the origin (n=31 colonies examined per construct). Eight is the median and the most common inferred minimum ploidy, consistent with previous estimates of ploidy in *E. coli*(10,23). Higher estimates of 16, 32, and 64 could have resulted from delayed ssDNA integration or ssDNA integration in a filamentous cell.

To further test this hypothesis, we inferred a minimal ploidy of the aforementioned target sites at the precise time point of ssDNA integration with an adapted *lacZ*-reporter assay that measured the fraction of mutant cells in mutant colonies directly after recombineering (Methods). Overall, the distributions of ploidy were similar for the three tested chromosomal locations at 5.5%, 9.7% and 34.2% genome distance from the origin (Fig. 3C). This result potentially explains why we observed no effect of chromosomal location on phenotypic delay. Furthermore, this principle should apply not only to recombineering, but also to natural mutations that arise during DNA replication.

### Effective polyploidy and phenotypic delay affect mutation rate estimation

Fluctuation tests are widely used to estimate bacterial mutation rates by counting mutants exhibiting a selectable phenotype. Selection is typically applied to stationary phase cells(22), which are expected to be polyploid(10,23). That polyploidy should affect the appearance of mutants in the fluctuation test was already pointed out quite early in the literature (24,25), but subsequently this consideration largely fell by the wayside.

When a mutation arises in an effectively polyploid cell, the first homozygous mutant descendant must appear as a single cell in the population (Fig. 4). Therefore in contrast to earlier interpretations(1), the frequent observation of singletons in the mutant distribution does not invalidate the existence of a substantial phenotypic delay. Furthermore, since heterozygous cells carrying recessive mutations do not exhibit the mutant phenotype, i.e. cannot form colonies on selective plates in the fluctuation test, these unobserved mutants should result in an underestimation of the mutation rate. Incidentally, the phage or antibiotic resistance mutations typically used in fluctuation tests are recessive(11,12,26).

**Fig. 4.**
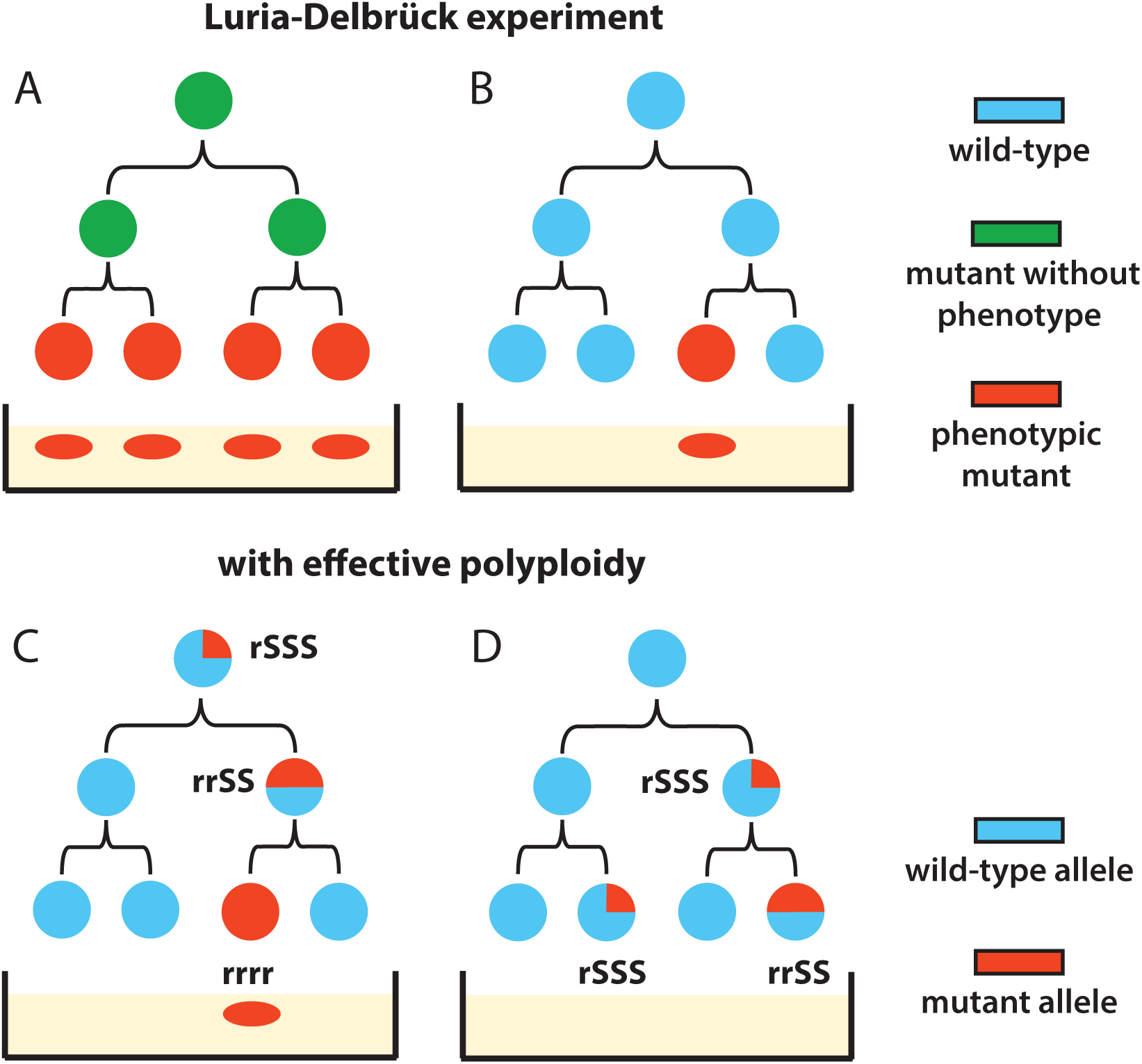
Reconciling Luria-Delbrück fluctuation test with phenotypic delay by effective polyploidy. (A): The original Luria-Delbrück mutation model disregards polyploidy. For instance, a phenotypic delay of 2 generations results in 4 mutants appearing at once. (B): The observation of many one-mutant (“singleton”) populations was interpreted as evidence against the existence of a delay(1,3). (C): With polyploidy considered, cells with four genome copies require two divisions to generate a homozygous mutant that expresses a selectable recessive phenotype. Hence a delay of two generations can generate just one mutant. (D): Heterozygous cells containing recessive mutations will not survive selection, leading to an underestimation of mutational events.

The impact of polyploidy on mutation rate estimation from fluctuation tests using the modern “gold standard” maximum likelihood method(22) has not been examined. Although one recent study considered the “segregation lag” for recessive mutations resulting from polyploidy(27), corrections to mutation rate estimators were only derived for two simpler methods with limited range of accuracy and low statistical efficiency relative to the maximum likelihood method(22). Furthermore, these derivations neglected the key point that not all descendants of heterozygous mutants will be mutants themselves.

We investigated the effect of polyploidy on observed mutant distributions and hence estimated mutation rates, for both dominant and recessive mutations, by simulating fluctuation tests. Our simulation model assumed fixed effective ploidy at the target, by doubling and symmetrically dividing chromosome copies upon division according to a model of segregation that is justified for *E. coli*(Methods). Importantly, this model leads to the shortest possible time to homozygosity, and thus a conservative estimate of lag; however, other models of segregation in bacteria and archaea are possible(27,28). From simulated cultures we counted phenotypic mutants given either a completely dominant or a completely recessive mutation, assuming instant protein equilibration, and estimated mutation rates using standard maximum likelihood methods(22,29,30). Other than polyploidy and dominance considerations, all modelling assumptions are the same as the standard approach (Methods), allowing us to isolate the effect of ploidy. Under our model, ploidy level *c* has two effects relative to monoploidy: (i) it increases the number of mutation targets and thus the per-cell mutation rate by a factor *c*, and (ii) it generates initially heterozygous mutants that after a delay of log2*c* generations produce one out of *c* homozygous mutant descendants (Fig. S1).

The mutation rate estimate at the mutational target can be compared to the actual per-copy rate µ_c_ and per-cell rate *c* µ_c_ used in the simulations (Fig. 5A-B). When *c*=1 (monoploidy), as the standard method assumes, the estimate indeed reflects the per-copy or (equivalently) per-cell rate. For *c*>1 the estimate is higher for dominant than for recessive traits. Surprisingly, for recessive traits, the estimate tends to coincide with the per-copy rate µ_c_ regardless of ploidy. For dominant traits, the estimate lies between the per-copy and per-cell rates, with confidence interval size increasing with ploidy. These patterns are robust across a range of parameter values (Fig. S2). In fact, these effects have a precise mathematical explanation (Suppl. Text section 2): the distribution of mutant counts in a polyploid population turns out to match the standard (monoploid) model with rescaled mutational influx in the case of a recessive trait (Fig. 5C and S3 A), but fundamentally differs for a dominant trait (Fig. 5D and S3B). Thus, for more commonly used recessive traits, estimates reflect mutation rates per target copy, which can be scaled up to per genome copy. This could explain why per-nucleotide mutation rates estimated from different targets do not differ significantly, despite differences in target location that potentially influence their copy number(31).

**Fig. 5.**
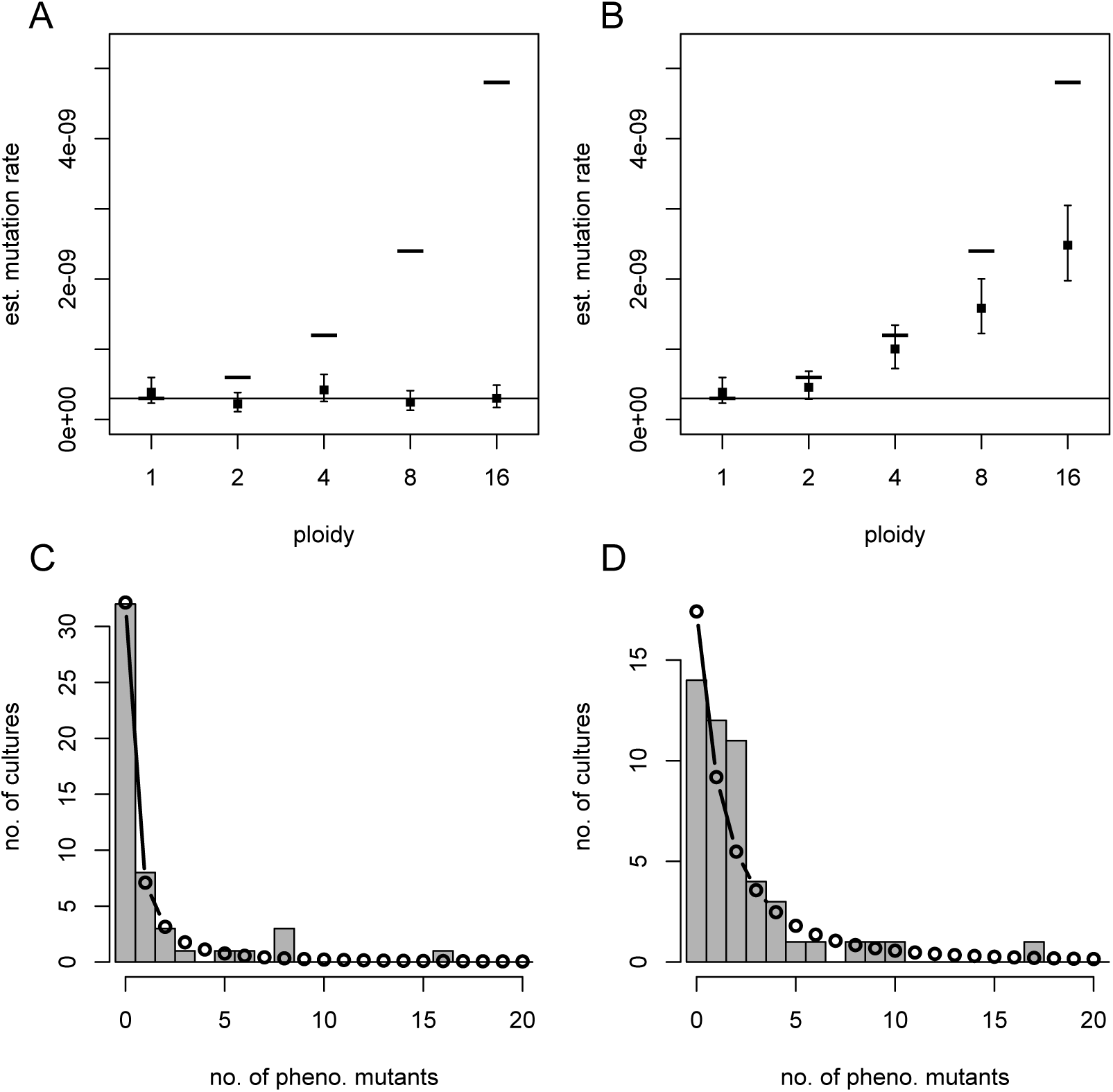
Simulated fluctuation tests. Mutation rate estimates (maximum likelihoodestimate, MLE, filled squares; and 95% confidence intervals, error bars) from 50simulated parallel cultures at each ploidy (c), with constant mutant interdivision time,assuming either a recessive (A) or dominant (B) mutation. The lower solid line and upper dashes indicate the per-copy (μ_c_ = 3 x 10^-10^) and per-cell (*c*μ_c_) mutation rates,respectively, used for simulation. For *c*=4 and recessive (C) or dominant (D) mutations,the observed mutant count distribution (histogram) is compared to that predicted by the standard model parameterized by the MLE mutation rate (connected points)

We then asked whether effective polyploidy impacts mutation rate estimates based on whole-genome sequencing (WGS) as well as fluctuation tests. WGS is typically conducted on evolved populations from mutation accumulation assays (MA) which use single-cell bottlenecking to minimize selection(31). Under the simplifying assumptions of fixed generation time and no cell death, we modeled an MA assay by tracking the single lineage that passes through each bottleneck and is ultimately sampled for sequencing (Suppl. section 3 and Fig. S4). Accounting for polyploidy, the per-cell mutation rate is *c*µ_g_ where µ_g_ is the per-genome-copy mutation rate. However, due to asymmetric inheritance, only a fraction 1/*c* of descendants from a mutant progenitor will eventually become homozygous mutants. Thus, only a fraction 1/*c* of mutations arising in the focal lineage will ultimately be sampled, leaving the per-genome copy rate µg as the inferred mutation rate (Suppl. section 3). We therefore conclude that neither the fluctuation test nor WGS methods can accurately capture the per-cell mutation rate. Thus, neglecting polyploidy underestimates the total influx of *de novo* mutations in a population, which is relevant for adaptation.

### Polyploidy and phenotypic delay impact bacterial evolvability under selection

Effective polyploidy has important consequences for evolutionary adaptation, both through the aforementioned increased influx of mutations and the masking of recessive mutations phenotype. Masking of deleterious recessive mutations is expected to increase their frequency in the standing genetic variation (SGV) and yield transiently lower, but eventually higher, mutational load in a fixed environment(2 8,3 2). This higher standing frequency could promote adaptation to new environments should these mutations become beneficial. However, in an environment where mutations are beneficial, masking their effects should hinder adaptation. Previous theoretical studies addressing these conflicting effects of ploidy on adaptation(3 2,3 3) have not been linked to bacteria, nor have they specifically considered the chance of evolutionary rescue, i.e. rapid adaptation preventing extinction under sudden harsh environmental change (e.g. antibiotic treatment).

Rescue mutations may pre-exist in the SGV and/or arise *de novo* after the environmental shift during residual divisions of wild-type cells. The source of rescue mutations has implications for the optimal approach to drug treatment(34) and the preservation of genetic diversity following rescue(3 5). To address the impact of effective polyploidy on rescue from SGV and *de novo* mutations, we developed a mathematical model of replication, mutations, and chromosome segregation in polyploid bacterial cells (Fig. 6 and Suppl. section 4).

**Fig. 6.**
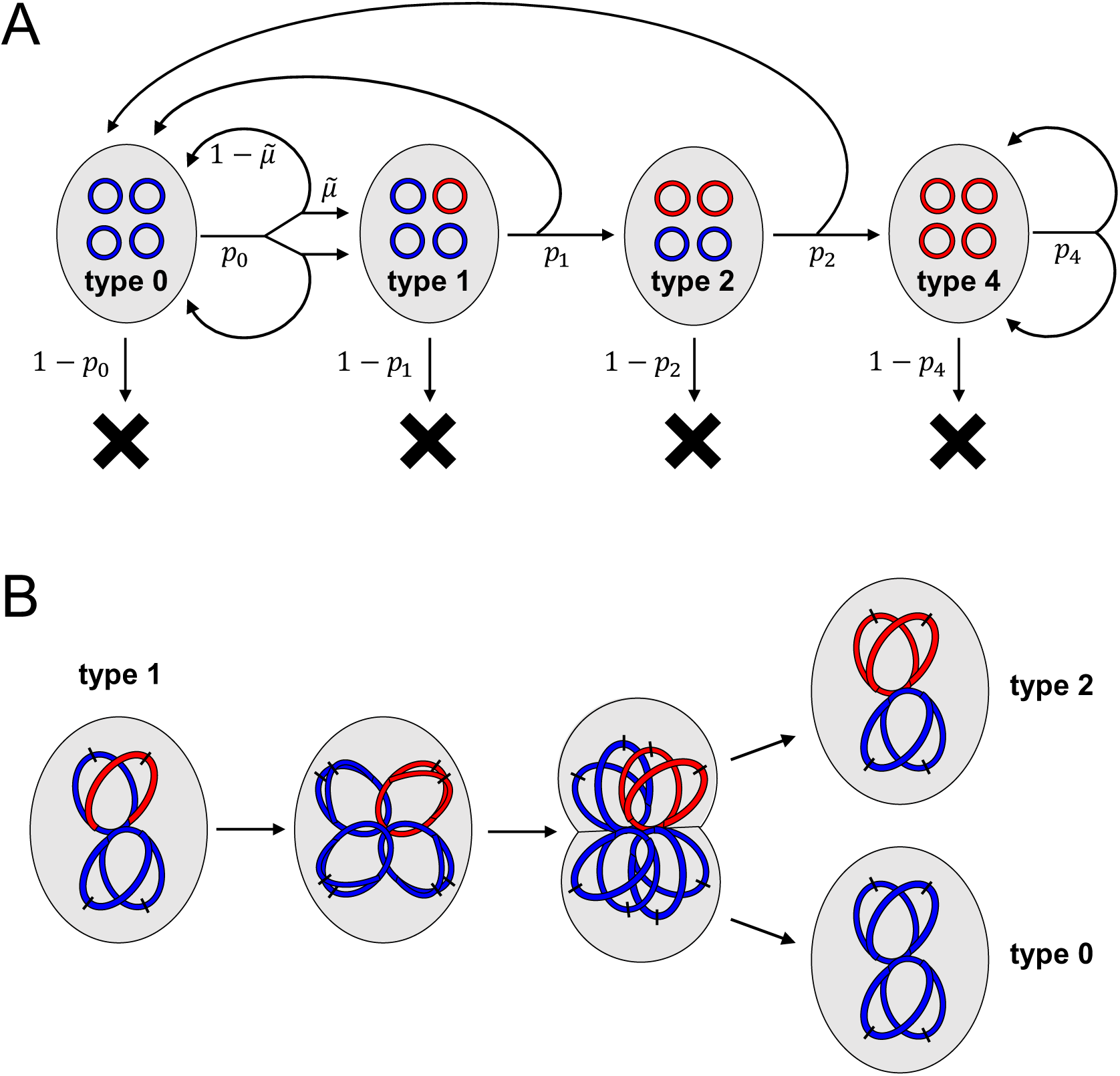
Schematic of the model used to evaluate standing genetic variation and rescue, illustrated for ploidy *c*=4. (A]: Flow diagram of all events. A cell is represented by a grey oval, containing four chromosomes (complete or partial, so long as they contain the gene of interest]. These chromosomes are coloured blue if wild-type at the gene of interest or red if mutant. A cell either divides to produce two daughter cells with type-specific probability *p*_*j*_, or otherwise dies. These probabilities *p_j_* differ between the old environment (to model SGV] and the new environment (to model rescue]. Upon type 0 division, mutation (producing type 1] occurs with probability 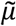in each daughter cell; otherwise the daughter is also type 0. In the remaining types, chromosome segregation determines the types of the daughter cells. (B]: A mechanistic view of chromosome replication and segregation, illustrated for the production of one type *2* and one type 0 daughter cell from a type 1 mother cell. On each chromosome, the black dash indicates the origin of replication.

We first derived the frequency of mutants in the SGV at mutation-selection balance in the “old” environment, where mutations continually arise at rate 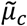 per chromosome copy per replication and, if expressed, carry a fitness cost *s* (Suppl. section 4.2). This yielded analytical expressions confirming that polyploidy increases the total frequency of a recessive mutant allele by masking its cost in heterozygotes. In contrast, the total mutant allele frequency is independent of ploidy if the mutation is dominant (Table 1 and Fig. S5).

**Table 1.** Approximate mutant frequencies at mutation-selection balance. Ploidy is *c* = *2*^n^ (*n* ≥ 1) in the polyploid cases; 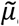_*c*_ is the per-copy mutation rate; and *s* is the cost of the mutation in homozygotes (in, the cost is masked in the recessive case but expressed in the dominant case).

|  | Monoploid | Polyploid<br>recessive | Polyploid<br>dominant |
| --- | --- | --- | --- |
| **Freq. of heterozyg. mutant cells**<br>($2^i$ mutant chroms., $0 \leq i \leq n-1$) | – | $2^{n-i} \tilde{\mu}_c$ | $(1-s)^i 2^{n-i} \tilde{\mu}_c$ |
| **Freq. of homozyg. mutant cells**<br>($2^n = c$ mutant chroms.) | $\tilde{\mu}_c/s$ | $\tilde{\mu}_c/s$ | $(1-s)^n \tilde{\mu}_c/s$ |
| **Total freq. of mutant allele** | $\tilde{\mu}_c/s$ | $(n-1) \tilde{\mu}_c + \tilde{\mu}_c/s$ | $\tilde{\mu}_c/s$ |

Next, we considered the fate of the population upon shifting to a new, harsh environment (e.g. antibiotic treatment), where phenotypically wild-type (“sensitive”) cells have a low probability of successfully dividing, while phenotypically mutant (“resistant”) cells have a higher probability. The population may already contain heterozygous and homozygous mutants in the SGV, and additionally gives rise to *de novo* mutants stochastically according to a Poisson process. We developed a branching process model to evaluate the probability that such mutations escape stochastic extinction, accounting in particular for the multiple cell divisions required until mutations segregate to homozygosity, with probabilities of successful division depending on whether these mutations are recessive or dominant. Finally, combining these model components yielded expressions for the probability of population rescue from SGV, *P*_SGV_, and from *de novo* mutations, *P*_DN_ (Suppl. section 4.3).

These rescue probabilities depend strongly on ploidy, dominance, and other model parameters (Fig. 7 and S6-7). In the recessive case, if phenotypically wild-type cells cannot divide in the new environment (e.g. a perfectly effective antibiotic), then *P*_SGV_ is independent of ploidy, reflecting the constant frequency of pre-existing phenotypically mutant homozygotes (Table 1). This result is consistent with our above findings for fluctuation tests. If division of wild-type cells is possible (e.g. imperfect antibiotic efficacy) then *P*_SGV_ increases with ploidy, since heterozygotes may produce additional homozygous mutant descendants. On the other hand, *P*_DN_ decreases with ploidy, as *de novo* mutations require more cell divisions until segregation is complete and the mutant phenotype is expressed, which turns out to outweigh the increase in mutational influx (Suppl. Text). Thus, although the overall probability of rescue remains similar as ploidy increases, rescue is increasingly from SGV rather than *de novo* mutations (Fig. 7A-B). These qualitative patterns are robust to variations in the model parameters (Fig. S6-7). For dominant mutations, on the other hand, both *P*_SGV_ and *P*_DN_ increase with ploidy, and their relative contributions can show more complex patterns (Fig. 7C-D and S6-7). In general, SGV makes a relatively larger contribution when the mutation has a low cost in antibiotic-free conditions (small *s*) and when the antibiotic is highly effective (low *p_S_*), in agreement with previous findings in the evolutionary rescue literature (36).

**Fig. 7.**
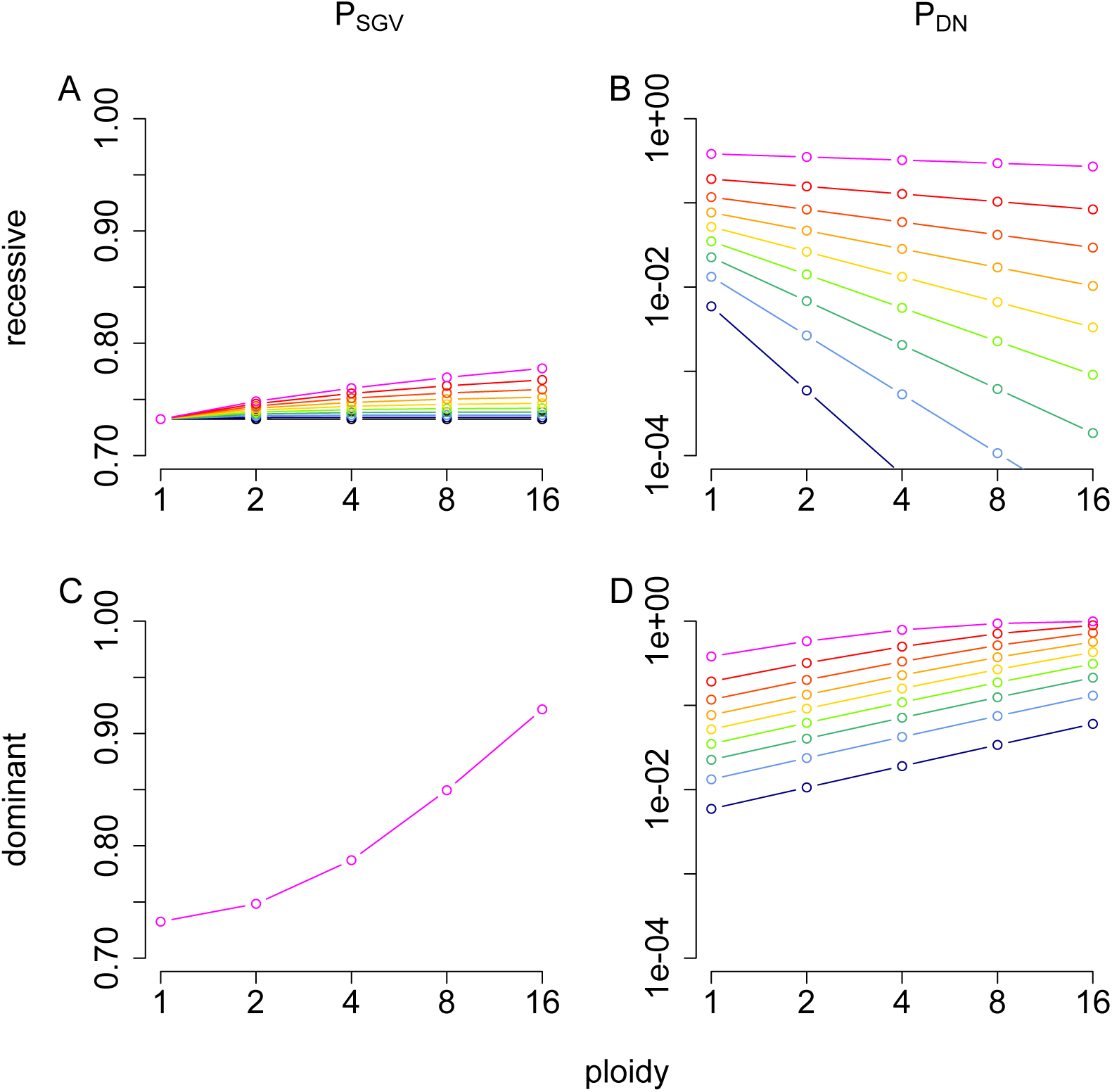
Impact of effective polyploidy on the probability of evolutionary rescue. (A,C): The probability that at least one mutation from the SGV survives in the new environment (*P*_SGV_; linear scale), and (B,D): the probability that at least one mutation arises in the new environment and survives (PDN; log scale), plotted as a function of ploidy, for a recessive mutation (A,B) or a dominant mutation (C,D). The different coloured curves indicate probability of division before death of phenotypically wild type(sensitive) cells in the new environment, *p*_S_, varying from 0 (black) to 0.45 (magenta) in increments of 0.05. The remaining parameters are fixed: probability of division before death of phenotypically mutant (resistant) cells, *p*_R_= 0.9; mutational cost in the old environment, *s* = 0.1; population size at the time the environment changes, *N* = 2 x 108; and per-copy mutation rate, 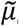 = 3 x 10-10.

## Discussion

The phenotypic effect of a bacterial mutation cannot manifest instantaneously. Here we thus asked two questions: how large is this phenotypic delay and what is its primary cause? We found a delay of 3-4 generations in the expression of three recessive antibiotic resistance mutations in *E. coli* and provided evidence that effective polyploidy is its primary cause.

Polyploidy is often regarded as a transient property limited to fast-growing bacteria, but this view has been challenged in recent years. Though ploidy tends to be higher during exponential growth (up to eight or 16 partial chromosome copies) (9), even during stationary phase *E. coli* cells contain typically four and up to eight complete chromosome copies(10). Environmental stresses can also induce multinucleated, polyploid cell filaments(37), in which adaptive mutations must overcome phenotypic delay before allowing population survival in deteriorating environments. A recent study exposing bacteria to low doses of the antibiotic ciprofloxacin showed that resistant bacteria can only emerge from mono-nucleated offspring cells that bud off from a long multinucleated cellular filament(37). This observation can be explained by masking of the mutant phenotype in polyploid, heterozygous cells. Furthermore, obligate polyploid bacterial species ranging from free-living bacteria to clinically relevant pathogens have been discovered across six phyla(23,38,39). This has for instance been recognized as a confounding factor in metagenomic studies of bacterial community structure by marker-gene-based analysis(38). Even within the same bacteria species, ploidy may vary in response to selection, as shown in a previous study that *E.coli* with resistance to camphor vapor also showed increased ploidy(40). Therefore, we argue that polyploidy is broadly relevant for bacteria, and will generally result in phenotypic delay of recessive mutations.

Dominance and polyploidy (whether effective or obligate) together affect the number of mutants observed in fluctuation tests and thus require reinterpretation of mutation rate estimates. Fluctuation tests typically use recessive antibiotic resistance mutations. Encouragingly, we found that the resulting estimates accurately reflect the per-target-copy mutation rate, regardless of ploidy. Therefore, studies using fluctuation tests to compare per-target-copy mutation rates across different conditions, e.g. (41), remain valid. Similarly, we showed that sequencing-based methods of mutation rate estimation from mutation accumulation assays reflect per-genome-copy rates. Thus, effective polyploidy does not appear to explain the up to ten-fold difference in mutation rate estimates (31,42) obtained using these two different methods.

Importantly, however, neither method reflects the per-cell mutation rate, and thus the total mutational influx in the population, which is proportional to ploidy. Indeed, our models suggest that fluctuation tests with recessive mutations or sequencing-based methods leave no detectable signal of ploidy in the data: that is, a polyploid population is indistinguishable from a monoploid population in these assays, even though their total mutational influx differs by a factor equal to the ploidy. Meanwhile, dominant mutations lead to fundamentally different mutant distributions in the fluctuation test, and neither the per-copy nor the per-cell mutation rate is accurately estimated. In conclusion, mutation rate estimates must be interpreted with caution, regardless of the method used.

The effects of polyploidy on number of mutational targets and phenotypic delay influence the evolutionary potential of populations to escape extinction under sudden environmental change such as antibiotic treatment. In particular, we showed that recessive rescue mutations are increasingly likely to come from the standing genetic variation as ploidy increases. This is due to the dual effects of masking the fitness cost of these mutations in the old (antibiotic-free) environment, while decreasing the chance that *de novo* mutations survive in the new (antibiotic) environment until their beneficial phenotype is expressed. Our novel results for rescue are broadly in line with previous theoretical findings on the role of ploidy for adaptation(32), and highlight the point that these considerations are relevant to bacteria as well as eukaryotes.

Our theoretical results rest on several simplifying assumptions. Firstly, we examined only the cases of complete dominance or complete recessivity. More generally, gene dosage effects could result in intermediate dominance; in this case, we expect the effects on mutation rate estimation and rescue probability to be intermediate between the two extremes considered here. Secondly, while we examined rescue via single mutations, if multiple mutations with different dominance were available, populations at different ploidy levels may tend to evolve via different pathways(43). Furthermore, while we exclusively considered chromosomal mutations, mutations on plasmids, particularly those with high copy number(44), should show similar effects, although segregation patterns and hence time to achieve homozygosity are likely to differ. Finally, models developed thus far have assumed constant ploidy, whereas future modeling efforts could incorporate the dynamically changing and environment-dependent nature of bacterial ploidy.

Given the manifold implications of a multigenerational phenotypic delay, we argue that effective polyploidy and the resulting phenotypic delay are essential factors to consider in future studies of bacterial mutation and adaptation.

## Materials and Methods

### Bacterial strains, antibiotics and media

All experiments were performed with strains derived from the wild-type *E. coli* MG1655 strain. A complete list of strains can be found in Table S2. Cells were grown at 30°C in LB or in M9 media with 0.4% lactose. Antibiotics were purchased from Sigma-Aldrich. To prepare stocks, rifampicin was dissolved in DMSO to 100 mg/ml; nalidixic acid was dissolved in 0.3M NaOH solution to 30 mg/ml; streptomycin and ampicillin were dissolved in MilliQ water to 100 mg/ml and filter sterilized. Rifampicin, streptomycin and ampicillin stocks were kept at −20°C while nalidixic acid was kept at 4°C. 100 mg/L ampicillin was used for maintaining the pSIM6 recombineering plasmid. All antibiotic agar plates were prepared fresh before every experiment.

### MIC determination

The MICs of rifampicin, streptomycin and nalidixic acid were determined by broth dilution method in LB and found to be 12 mg/L, 12 mg/L and 6 mg/L, respectively.

### High efficiency recombineering

Our recombineering protocol was adapted from previous studies(6,21). To ensure reproducibility, a detailed step-by-step protocol is provided in the Supplementary Text (Section 1). In brief, *E. coli* harboring pSIM6 plasmids were grown into early exponential phase before heat-activation at 43°C for 10 minutes to express the recombineering proteins. Activated cells were then repeatedly washed in ice-cold MilliQ water to remove residual salts. 50 µl of concentrated salt-free cell suspension was then mixed with ~200 ng of ssDNA before electroporation at 1.8 kV/mm. Immediately after electroporation cells were resuspended in LB and recovered for 30 min at 30°C. After this initial recovery cells were pelleted, then resuspended in fresh LB to continuously grow at 30°C for subsequent phenotyping.

### Quantification of phenotypic delay of resistance mutations

From the resuspended population ~2% of cells were sampled hourly for the first 10 hours and then at 24 hours. A time point at 48 hours was also included, to control for factors that potentially prevent phenotypic penetrance from ever reaching 100%, such as low establishment probability of mutant cells. The sampled populations were appropriately diluted for optimal plating onto selective and non-selective plates. Total population size and thus generations elapsed in the sampled cultures was estimated from CFU on non-selective plates. To score the frequency of genotypic mutants we replica-plated all colonies from the non-selective plates to selective plates for each tested time point. The frequency of genotypic mutants, *F*_*g*_, was determined by the fraction of colonies from non-selective plates that could grow after postponed replica-plating onto selective plates. The frequency of phenotypic mutants, *F*_*p*_, was determined by the ratio of CFU from immediate plating on selective plates versus CFU on non-selective plates. Phenotypic penetrance was defined as *P* = F_*p*_/ F_*g*_.Phenotypic delay was then quantified as the time point at which phenotypic penetrance reaches 50%.

### Quantification of homozygosity

To quantify mutant homozygosity, i.e. the fraction of homozygous mutants among all genotypic mutants, we developed a *lacZ*-based visual assay. We constructed bacterial strains with a *lacZ* gene disrupted by a nonsense point mutation (E461X)(8) and inserted the broken *lacZ* within 5 kb of each antibiotic resistance target gene. These strains were subjected to recombineering with an ssDNA carrying the reverse point mutation (X461E) that restored the lac+ phenotype. The resulting phenotypic mutants were selected on M9-lactose media. Phenotypic mutants become blue on permissive media containing 1 mM IPTG and 40 μg/ml X-gal(5). Heterozygous mutants with mixed lac+/lac alleles form blue/white-sectored colonies, whereas homozygous mutants form entirely blue colonies (Fig. 2E). Plates with colonies were left at 4°C for one week to allow sufficient development of the blue color, but before the blue pigment spreads too far to obscure sectored colonies. Counting sectored (*s*) and non-sectored (*n*) blue colonies, we determined mutant homozygosity as *f*_hom_ = *n*/(*s*+*n*). Comparing *f*_hom_ to the phenotypic penetrance *P* thus indicates to what extent phenotypic delay is attributable to effective polyploidy. Colony counting was performed using CellProfiler(45).

### Single-cell observations

We constructed a strain with a constitutively expressed *YFP* gene disrupted by three consecutive stop codons. Recombineering corrected the stop codons. After electroporation and 30 min recovery at 30°C, 1 μl of appropriately diluted cell suspension was pipetted onto a small 1.5% UltraPure Low Melting Point agarose pad. After drying the pad for 1 minute it was deposited upside down in a sealed glass bottom dish (WillCo Wells, GWST-5040). Time-lapse microscopy was performed with a fully automated Olympus IX81 inverted microscope, with 100X NA1.3 oil objective and Hamamatsu ORCA-flash 4.0 sCMOS camera. For fluorescent imaging, we used a Lumen Dynamics X-Cite120 lamp and Chroma YFP fluorescent filter (NA1028). The sample was maintained at 30°C by a microscope incubator. Phase-contrast and yellow fluorescence images were captured at 5-minute intervals for 16 hours. The subsequent image analysis was performed with a custom-made MATLAB program (Vanellus, accessible at: http://kiviet.com/research/vanellus.php)

### Assessing minimal ploidy

We performed the *lacZ*-reporter assay, as described above for three strains with *lacZ*-gene juxtaposed to each of the antibiotic resistance target genes. After the 30-min recovery following recombineering (before extensive growth), cells were plated directly onto LB agar with IPTG and X-gal. After 24 hours of incubation at 30°C, entire mutant colonies that contained blue color were picked. We started from colonies closest to the center of each agar plate and expanded outwards to eliminate picking bias. The picked colonies were diluted 104- to 105-fold in PBS before plating on average about 500 CFUs on fresh LB agar with IPTG and X-gal to infer the fraction of mutant cells in the given colony. This fraction was then used to deduce the minimal ploidy at the time of ssDNA integration based on a previous study(46): a colony with one-quarter mutant cells for example has minimal ploidy of two, as it could have resulted from mutagenesis on one out of four DNA single-strands. Actual ploidy may be higher, if for example two out of eight single strands mutated in a cell of ploidy four.

### Ploidy and chromosome segregation model

For simplicity, we assumed every cell has the same effective ploidy, i.e. copies of the gene of interest, over the relevant timescale. At each generation, chromosomes must thus undergo one round of replication and be evenly divided between the two daughter cells. In *E. coli*, chromosomes appear to progressively separate as they are replicated and detach last at the terminus(9). We therefore assumed segregation into daughter cells occurs at the most ancestral split in the chromosome genealogy. This assumption is conservative because it implies that mutant chromosomes always remain together, resulting in the fastest possible approach to homozygosity and thus the shortest phenotypic delay. Under this model, ploidy must take the form *c* = 2^*n*^ (for *n* = 0, 1, 2,…), among which the number of mutant copies is *j* = 0 or 2*i*(0 ≤ *i* ≤ *n*), while the remaining *c* -*j* copies are wild-type. Note that other models of segregation are possible, e.g. random segregation in highly polyploid Archaea(28), which would lead to slower approach to homozygosity and corresponding effects on the evolutionary model results.

### Simulated fluctuation tests

All simulations and inference were implemented in R. We wrote our own code to account for polyploidy, but in the future our methods could potentially be integrated into recently published R packages for fluctuation analysis (47,48). We simulated culture growth in non-selective media with stochastic appearance of spontaneous *de novo* mutations (for details see Supplementary Text, Section 3.1). We assumed a fixed per-copy mutation rate of μ_*c*_ per wild-type cell division, such that the per-cell mutation rate is μ = *c* μ_*c*_ for effective ploidy *c*. We neglected the chance of more than one copy mutating simultaneously, i.e. mutants always arose with the mutation in a single chromosome copy. Furthermore, we only counted mutations once they arose in double-stranded form. The descendants of each *de novo* mutant were tracked individually, with mutant chromosomes segregating as described above and interdivision times either drawn independently from an exponential distribution or constant. We assumed no fitness differences between wild-type and mutant in nonselective media. In the case of *c*=1 and exponential interdivision times our model corresponds to the standard “Lea-Coulson” model(22,30), which is also the basis of the widely used software FALCOR(49).

Each simulated culture was initiated with 1000 wild-type cells, and after 20 wild-type population doublings, the culture growth phase ended and phenotypic mutants were counted, under the assumption of either complete recessivity (requiring all *c* chromosomes to be mutant) or complete dominance (requiring at least one mutant chromosome). Assuming (as standard) 100% plating efficiency and no growth of phenotypically wild-type cells under selective conditions, the number of colonies formed on selective plates equals the number of phenotypic mutants in the final culture. The mutant colony counts from 50 simulated parallel cultures were then used to obtain a maximum likelihood estimate 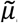 and 95% profile likelihood confidence intervals of mutation rate under the standard model, which in particular assumes that a *de novo* mutant and all its descendants are immediately phenotypically mutant. The best-fitting distribution of mutant counts was calculated from the standard model with mutation rate equal to 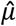. While we implemented these calculations in R, calculation of the likelihood under this model has been previously described(29,50) and has also been implemented in FALCOR(49).

### Mutation-selection balance

We considered a population with effective ploidy *c*, in which mutations arise (again, in double-stranded form) in a proportion 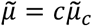 of offspring in each generation. The definition of mutation rate used in the population genetics literature is subtly different from that used in fluctuation analysis and thus given different notation here (see Supplementary Text, section 4.2). The mutation has relative fitness cost *s* in homozygotes, with the cost either completely masked (if recessive) or equal (if dominant) in heterozygotes. We extended deterministic genotype frequency recursions to incorporate chromosome segregation as described above, and solved for the equilibrium frequencies of all heterozygous and homozygous mutant types (Supplementary Text, section 4.2).

### Evolutionary rescue

We modeled the fate of a population shifted to a harsh new environment, i.e. either extinction or rescue by mutants, stochastically using a multi-type branching process. Unlike in the fluctuation test simulations, where we neglected the chance that wild-type cells produce surviving lineages in the new environment, here we allowed a probability *p*_S_ ≤ ½ that a phenotypically wild-type cell successfully divides before death to produce two offspring, while phenotypically mutant cells have corresponding probability *p*_R_ ≤ *½*. Thus phenotypically wild-type cells cannot sustain themselves, but have a non-zero chance of producing phenotypically mutant descendants either by segregation of mutant alleles in the standing genetic variation (SGV; modeled by mutation-selection balance as above), or *de novo* mutations during residual divisions in the new environment. We derived analytical approximations (Supplementary Text, section 4.3) for the probability of rescue from SGV (*P*_SGV_) or from*de novo* mutations (*P*_DN_), which are not mutually exclusive.

## Acknowledgements

We thank Erik Wistrand-Yuen, Erik Lundin and Gerrit Brandis from the groups of Dan I. Andersson and Diarmaid Hughes at Uppsala University for kindly gifting us some bacterial strains and plasmids. We also thank Sébastien Wielgoss, Gabriel Leventhal, Andrew Read and Min Wu for helpful discussions.

## Financial disclosure

This work was supported by the ETH Zurich; the European Research Council under the 7th Framework Programme of the European Commission (PBDR: grant number 268540 to SB); and the Swiss National Science Foundation (grant number 155866 to SB and grant number 31003A_149267 to MA).

## Author Contributions

**Conceptualization:** Lei Sun, Helen K. Alexander, Martin Ackermann, Sebastian Bonhoeffer.

**Formal analysis:** Lei Sun, Balazs Bogos, Daniel J. Kiviet, Helen K. Alexander.

**Funding acquisition:** Martin Ackermann, Sebastian Bonhoeffer.

**Investigation:** Lei Sun, Balazs Bogos, Daniel J. Kiviet, Helen K. Alexander.

**Methodology:** Lei Sun, Balazs Bogos, Daniel J. Kiviet, Helen K. Alexander.

**Project administration:** Sebastian Bonhoeffer.

**Software:** Balazs Bogos, Daniel J. Kiviet, Helen K. Alexander.

**Supervision:** Martin Ackermann, Sebastian Bonhoeffer.

**Writing - original draft:** Lei Sun, Helen K. Alexander, Sebastian Bonhoeffer.

**Writing - review & editing:** Lei Sun, Helen K. Alexander, Balazs Bogos, Daniel J. Kiviet, Martin Ackermann, Sebastian Bonhoeffer.

